# Turning a Kv channel into hot and cold receptor by perturbing its electromechanical coupling

**DOI:** 10.1101/2024.08.08.607202

**Authors:** Bernardo I Pinto-Anwandter, Carlos A Z Bassetto, Ramon Latorre, Francisco Bezanilla

**Author notes:** These authors contributed equally to this work (Bernardo I Pinto-Anwandter, Carlos A Z Bassetto).

## Abstract

Voltage-dependent potassium channels (Kv) are extremely sensitive to membrane voltage and play a crucial role in membrane repolarization during action potentials. Kv channels undergo voltage-dependent transitions between closed states before opening. Despite all we have learned using electrophysiological methods and structural studies, we still lack a detailed picture of the energetics of the activation process. We show here that even a single mutation can drastically modify the temperature response of the *Shaker* Kv channel. Using rapid cell membrane temperature steps (Tsteps), we explored the effects of temperature on the ILT mutant (V369I, I372L, and S376T) and the I384N mutant. The ILT mutant produces a significant separation between the transitions of the voltage sensor domain (VSD) activation and the I384N uncouples its movement from the opening of the domain (PD). ILT and I384N respond to temperature in drastically different ways. In ILT, temperature facilitates the opening of the channel akin to a “hot” receptor, reflecting the temperature dependence of the voltage sensor’s last transition and facilitating VSD to PD coupling (electromechanical coupling). In I384N, temperature stabilizes the channel closed configuration analogous to a “cold” receptor. Since I384N drastically uncouples the VSD from the pore opening, we reveal the intrinsic temperature dependence of the PD itself. Here, we propose that the electromechanical coupling has either a “loose” or “tight” conformation. In the loose conformation, the movement of the VSD is necessary but not sufficient to efficiently propagate the electromechanical energy to the S6 gate. In the tight conformation the VSD activation is more effectively translated into the opening of the PD. This conformational switch can be tuned by temperature and modifications of the S4 and S4-S5 linker. Our results show that we can modulate the temperature dependence of Kv channels by affecting its electromechanical coupling.

## Introduction

Voltage-gated potassium channels (Kv) regulate neuronal excitability and muscle contraction (1). Kv channels are tetramers with subunits containing six transmembrane segments (S1-S6). The S1-S4 segments comprise the voltage sensor domain (VSD), and the S5, P-loop, and S6 segments form the pore domain (PD) (2). The S4-S5 linker connects the voltage sensor to the pore domain. This linker plays a crucial role in the transduction of the movement of the VSD upon changes in membrane voltage with the opening and closing of the PD gate (3, 4).

Temperature is an intensive physical property that affects essentially all biological processes. Temperature changes have been extensively used to study the properties of Kv channels (5–10). The *Shaker* K^+^ channel is a well-established prototypic Kv channel, and its biophysical characteristics were widely characterized using electrophysiological, spectroscopic, and, more recently, structural methods (11–13). Its temperature dependence has been explored in great detail, showing that the channel’s open probability (Po) does not have a substantial temperature dependence (5, 6). However, some transitions in the activation pathway have high enthalpic and positive entropic changes, indicating a significant temperature dependence (5).

We rationalized that the temperature dependence of the Po depends on the strict coupling between the VSD and PD, a hallmark of *Shaker*-like Kvs (12, 14). Thus, by altering the coupling between the VSD and the PD, we should be able to modify the energy landscape and, therefore, the temperature dependence of the Shaker channel activation pathway. To approach this experimentally, we used the S4 triple mutant V369I, I372L, and S376T (ILT) and the single mutant I384N, a residue in the S4-S5 linker. The ILT mutation induces a large voltage separation between the main charge movement and the final voltage-dependent step leading to channel opening (15, 16). In contrast, the I384N mutation uncouples the voltage sensor movement from the pore opening (17).

To dissect the energy landscape of activation in these channels, we used our recently developed framework for fast temperature changes and measurement at the cell membrane under voltage control (18). We used our ability to control temperature and voltage to determine their effects on the movement of the VSD (gating currents) and opening of the PD (ionic currents). The ILT mutant allowed us to determine the thermodynamics of the last transition of the voltage sensor. Specifically, temperature facilitates the last step of the VSD movement, increasing the Po of the ILT mutant, thus becoming a hot receptor. In contrast, temperature causes a substantial decrease in the Po of the I384N mutant in a process independent of the VSD movement, therefore revealing an intrinsic temperature dependence of the pore and producing a cold receptor.

Our study provides new insights into the mechanisms underlying ion channel function and the role of the S4 and S4-S5 linkers in the temperature dependence of Kv channels. Our results show that the temperature dependence of a channel arises from the intrinsic properties of the VSD, the PD, and the coupling between them. Moreover, Kv channels can be converted into cold or hot receptors using a handful of mutations.

## RESULTS

### Tsteps reveal hot and cold sensing properties of ILT and I384N

We measure the temperature dependence of ionic conduction using our recently developed framework to generate fast temperature steps on the oocyte membrane (**Fig 1A**, Methods)(18). Since previous studies have indicated that the last concerted step in the opening of the channel has a high-temperature dependence (5, 6), we used two mutants that uncouple the voltage sensor to the pore domain at different locations, namely the ILT and I384N. The ILT mutations are located next to the C-terminus of S4, while the I384N is located next to the N-terminus of the S4-S5 linker, where it interacts with the C-terminus of the S6 (**Fig. 1B, C**). Using the Tstep method, we can apply temperature steps at any time during the activation process. After the ionic currents reached the steady state, we applied a temperature step in the middle of a voltage pulse protocol and analyzed the effect on ionic currents. In Shaker (WT) as soon as the temperature changes, we observe an increase in the ionic current which is more pronounced at depolarized potentials (∼ 40% increase at +40 mV for 8°C Tstep) (**Fig. 1D**). For the ILT mutant, we observed a more pronounced increase in the current (∼ 100% increase at +180 mV for 5°C Tstep) (**Fig. 1E**) while in the I384N mutant temperature induced a decrease in the current (∼ 30% decrease at +150 mV for 7°C Tstep) (**Fig. 1F**). When the time course of currents and temperature at different voltages were analyzed in the WT, we observed that at positive potentials (+ 40 mV) the current followed the time course of temperature, however at less depolarized voltages (< 0 mV) it deviated and decreased after an initial increase (**Fig 1G**). The rapid increase in the current upon application of the Tstep is consistent with an increase in the single-channel conductance (Δγ), while the decrease in current is related to a reduction of the Po. For the ILT, we observed an increase in the current and Po (ΔP_o_) at all the voltages tested (**Fig. 1H**). In contrast, for I384N, the ionic current initially increased due to the single channel conductance and then decreased due to a change in Po for all voltages tested (**Fig. 1I**). To compare the effects of temperature between the WT and mutants, we calculated the temperature coefficient (Q_10_) of the steady state current after the Tstep (Eq 3). The Q_10_ for the WT saturates at 1.5 for voltages over 0 and decreases at voltages < 0 mV due to the Po changes with temperature (**Fig. 1J**). The maximum Q_10_ is close to the 1.44 value determined previously for the single-channel conductance (5). The Q_10_ of the current induced by the ILT mutant is significantly higher than that of the WT. If the Tsteps were applied when the bath temperature was 5.6 °C, the Q10 varies between 5-20 depending on the voltage. The results were more modest at higher bath temperatures (2-2.5 at 22 °C), likely due to the nonlinearity of Q_10_ with temperature (19, 20) (**Fig 1K**). For I384N, when the bath temperature was 6 °C, the Q_10_ varied between 0.8-1, whereas when the bath temperature was 17 °C the Q_10_ further decreased to 0.5-0.6 (**Fig. 1L**). Since we expect these changes in temperature dependence to be caused by the changes in VSD to PD coupling, we explored the effect of temperature on the VSD movement.

**Figure 1:**
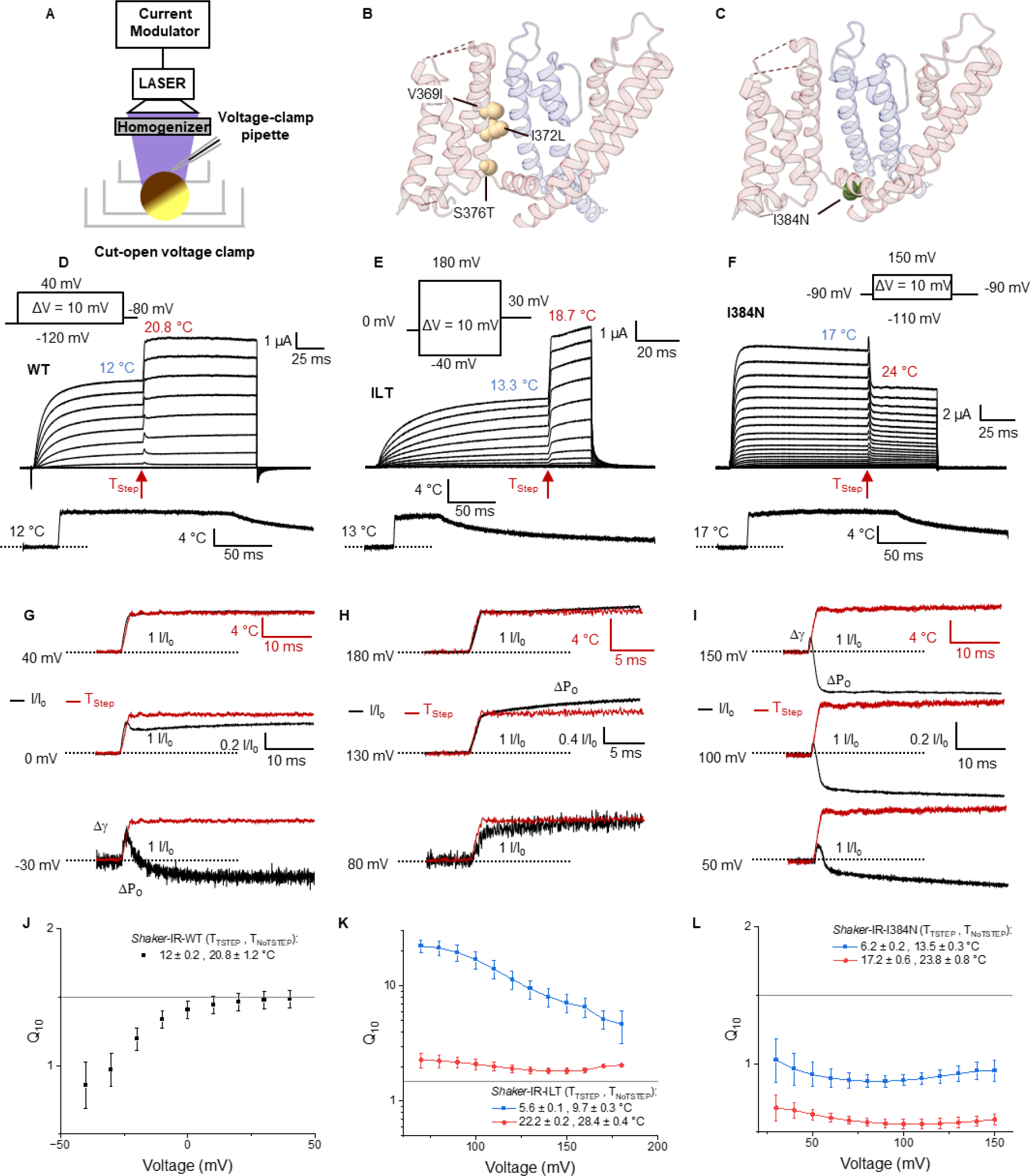
Temperature Dependence of Ionic Conduction in WT and Mutant Channels. **A)** Schematic representation of the experimental setup used to apply fast-temperature steps to the oocyte membrane. A current-regulated homogenized laser light is used to illuminate the upper dome of the oocyte in the cut-open setup, where the current recording occurs. The absorption of the visible light by melanin under the membrane of the oocyte generates a brief temperature change that is modulated by using PWM of the laser light. **B)** Structural location of the ILT mutations highlighting the location in the N-terminus of S4. **C)** Structural location of the I384N mutation at the C-terminus of the S4-S5 linker, indicating its interaction with the S6 segment in the pore domain (PD), interacting pore subunit shown in blue. **D-F)** Current response to temperature steps during a voltage pulse protocol for **(D)** Wild-type (WT) Shaker IR, **(E)** ILT and **(F)** I384N channels. **G-I)** Comparison of the time course of currents and temperature at different voltages for **(G)** WT, **(H)** ILT and **(I)** I384N channels. **J-L)** Plot of the Q10 of ionic currents for the **(J)** WT, **(K)** ILT and **(L)** I384N channels, colors indicate different initial bath temperatures.

### Temperature dependence of the voltage sensor

To determine the effect of temperature on the VSD movement, we measured the gating currents to obtain the charge movement. To characterize the behavior of the voltage sensor, we used the V478W mutant, which is known to render a non-conducting channel (21). We favor the use of this mutation over other mutations to measure gating currents, such as W434F, because of the absence of residual ionic current in this mutant (22, 23) and given that temperature changes induced ionic currents through the W434F pore (**Supp Fig. 1**). Similarly to experiments performed with ionic currents, we applied a Tstep in the middle of a voltage step after the gating currents had subsided and subtracted the linear optocapacitive component. We observe the development of inward currents from -120 to -30 mV in response to the Tstep (**Fig 2. A, B**). We can interpret the results on the gating currents as a change in the equilibrium distribution of the VSD through different conformational states with temperature. To verify this interpretation, we calculated the difference in the charge (ΔQ) displaced with and without Tsteps at different voltages. The charge difference between the temperature of 16°C and 22 °C shows an inverted bell shape curve with a minimum of around -50 mV (**Fig 2C**). This charge movement provides the underlying mechanism to explain the closure of the pore of the WT channels observed in the ionic currents (**Fig. 1D, G**). Since this only happens at specific voltage ranges, we only see the effect on the ionic currents at voltages where temperature displaces the VSD. Our analysis shows that Tsteps drive the VSD into the resting state in a time-dependent manner. Thus, we can predict that applying a Tstep at different times before a voltage step would produce a time-dependent delay in the activation because deeper closed states become populated. This mechanism would manifest in the classical Cole-Moore shift, which describes the delay in the opening due to the transitions of the VSD along closed states before the pore can open (24). To test this, we applied temperature steps at different times before the onset of a depolarizing voltage pulse (**Fig. 2D**). We observed that the kinetics of the opening in the presence of Tstep is faster when compared with the absence of Tstep as predicted. More importantly, the earlier the Tstep is applied before the test voltage step, the longer the delay due to the transition of the VSD into deeper close states (**Fig. 2 E,F**). As expected, the delay in the Cole-Moore observed in the ionic currents follows the time course of the charge movement with Tstep. These experiments, which can only be observed by precisely controlling temperature and voltage, corroborate our observation that the movement of the VSD by temperature produces the closure of the channel.

**Figure 2:**
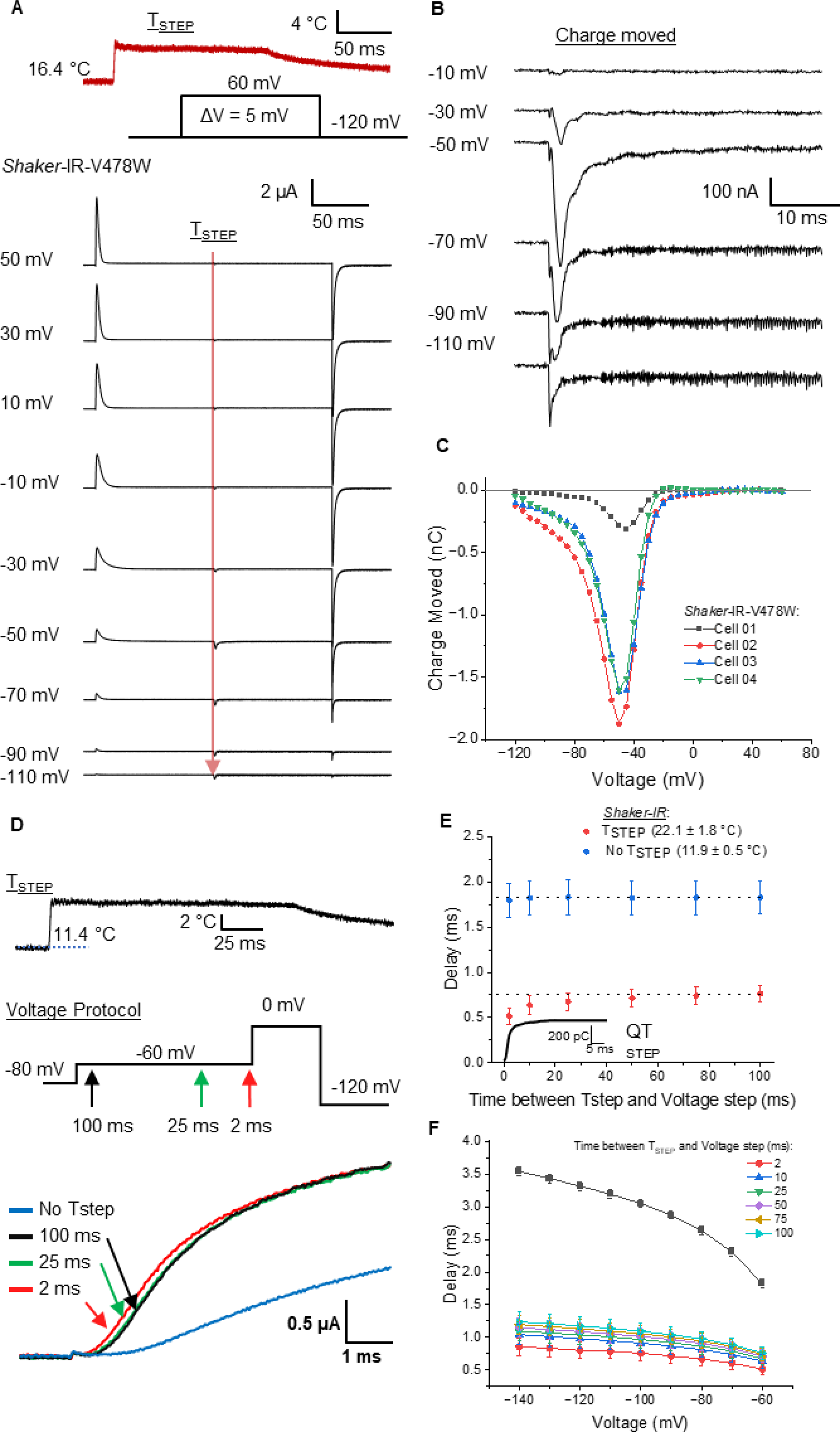
VSD displacement in response to temperature produces a time-dependent delay in activation. (**A)** Gating current measurements during a voltage and Tstep protocol. Temperature steps (Tsteps) were applied in the middle of a voltage step, after gating currents had subsided (indicated by the red arrow). **(B)** Detail of temperature-dependent charge movement. **(C)** Temperature-dependent charge movement vs voltage curves. **(D)** Ionic currents in response to a voltage step protocol, shown without a temperature step (No Tstep, blue) and with Tstep applied of 100 ms (black), 25 ms(green), and 2 ms (red) before the voltage pulse. The inset depicts the voltage protocol; arrows indicate time at which the Tstep is applied. Below, a trace representing the Tstep, note different time scales. **(E)** Delay in ionic current onset as a function of the time between Tstep and voltage step. Inset indicates the time course of the charge movement in response to a similar Tstep in V478W at -60 mV. **(F)** Relationship between delay in ionic current onset and voltage, for different Tstep durations applied before the test 0 mV voltage step (N=4 independent measurements, Temperatures =12° to 22° C).

### VSD displacement in response to temperature of ILT and I384N differs from the WT

Temperature promoted two distinct effects on the ILT gating currents. We observed an inward current from -100 mV to -50 mV and an outward current at voltages higher than 40 mV (**Figure 3A, B**). The displaced current shows a minimum at around -80 mV and a maximum at around +80mV (**Figure 3C**). This indicates that in the initial movement of the VSD, increasing temperature promotes a relative stabilization of the resting state, as it is in the WT channel. However, at more positive voltages (> 40 mV), temperature favors the voltage sensor active state, which will favor the opening of the pore producing a significant increase in the ionic current.

**Figure 3:**
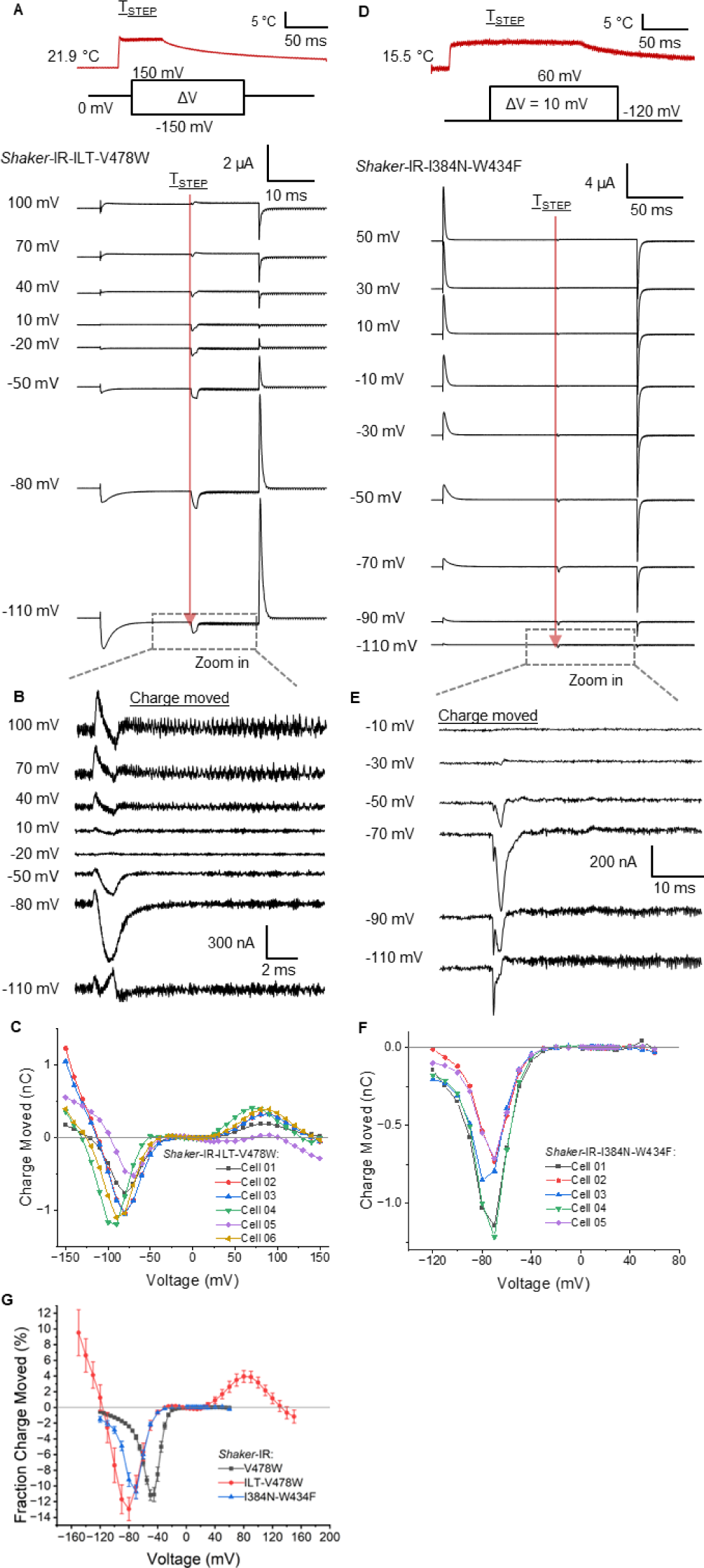
Temperature-dependent VSD displacement in ILT and I384N. (**A-B)** Gating current measurements during a voltage and Tstep protocol for **(A)** ILT-V478W and **(B)** I384N-W434F. Temperature steps (Tsteps) were applied in the middle of a voltage step after gating currents had subsided (indicated by the red arrow), to analyze the effect of temperature on gating charge movement. **(C, D)** Detail of temperature-dependent charge movement on **A, B**. (**E, F)** Temperature-dependent charge movement vs voltage curves for **(E)** ILT-V478W and **(F)** I384N-W434F. **(G)** Fraction of the total gating charge moved at different voltages for a WT= from 15.6 ± 0.3 °C to 22.8 ± 0.7 °C, ILT = from 22.2 ± 0.3 °C to 30.5 ± 0.9 °C, and I384N = from 15.4 ± 0.2 °C to 22.8 ± 1.1 °C step.

For I384N, due to the lack of expression when we used the mutation V478W, we used the W434F mutation to record the movement of gating charges. The W434F mutant presents a small conductance at voltages larger than -30 mV, a voltage range where the pore starts to open, this prevented further analysis using only this mutation on the channel. However, we still observed the inward current in response to a temperature jump indicative of VSD movement into the resting state, like V478W (**Suppl. Fig. 1**). We replaced the internal K^+^ from the oocytes, which allowed us to observe only gating currents in the I384N_W434F mutant (See methods). The Tstep in this mutant induced an inward current at voltages of -110 mV to -30 mV (**Figure 3D,E**). The displaced current shows a minimum around -70 mV (**Figure 3F**). At voltages larger than -30 mV, no charge displacement is observed, indicating that the decrease in the observed ionic currents is not associated with VSD movement. We can interpret the results on the gating currents as a change in the equilibrium distribution of different states of the VSD with temperature. When ΔQ is compared to the total gating charge, we find that Tsteps can move up to 10% of the total charge (**Fig. 3G**).

### A thermodynamic mechanism for temperature dependence

To interpret these results, we calculated the enthalpic (ΔH) and entropic (ΔS) components for both the conductance vs voltage (G-V) and charge vs voltage (Q-V) curves, using Tstep protocols that reproduce bath temperature changes (**Suppl. Fig. 2-4**). The G-V curves of the WT, ILT and I384N shift approximately 0.9, -1.7 and 2.4 mV/°C, respectively (**Fig. 4A-C**, **Table I**). Using the G-V measurements, we calculated the free energy of opening at different temperatures and obtained the ΔH and ΔS components (**Eq. 6**). For WT and I384N, ΔS is negative, while for ILT, it is positive, as expected from their respective shifts observed in the G-V curves (**Fig. 4D**, **Table I**).

**Table 1:** Thermodynamic parameters from G-V curves.

|  | **WT** | **ILT** | **I384N** |
| --- | --- | --- | --- |
| **Shift (mV/°C)** | 0.9±0.1 | -1.7±0.3 | 2.4±0.4 |
| **ΔH kcal/mol** | -17±8 | 10±2 | -22±5 |
| **ΔS cal/(mol\*K)** | -53±26 | 21±8 | -88±19 |

Values shown as Mean±SE

**Figure 4:**
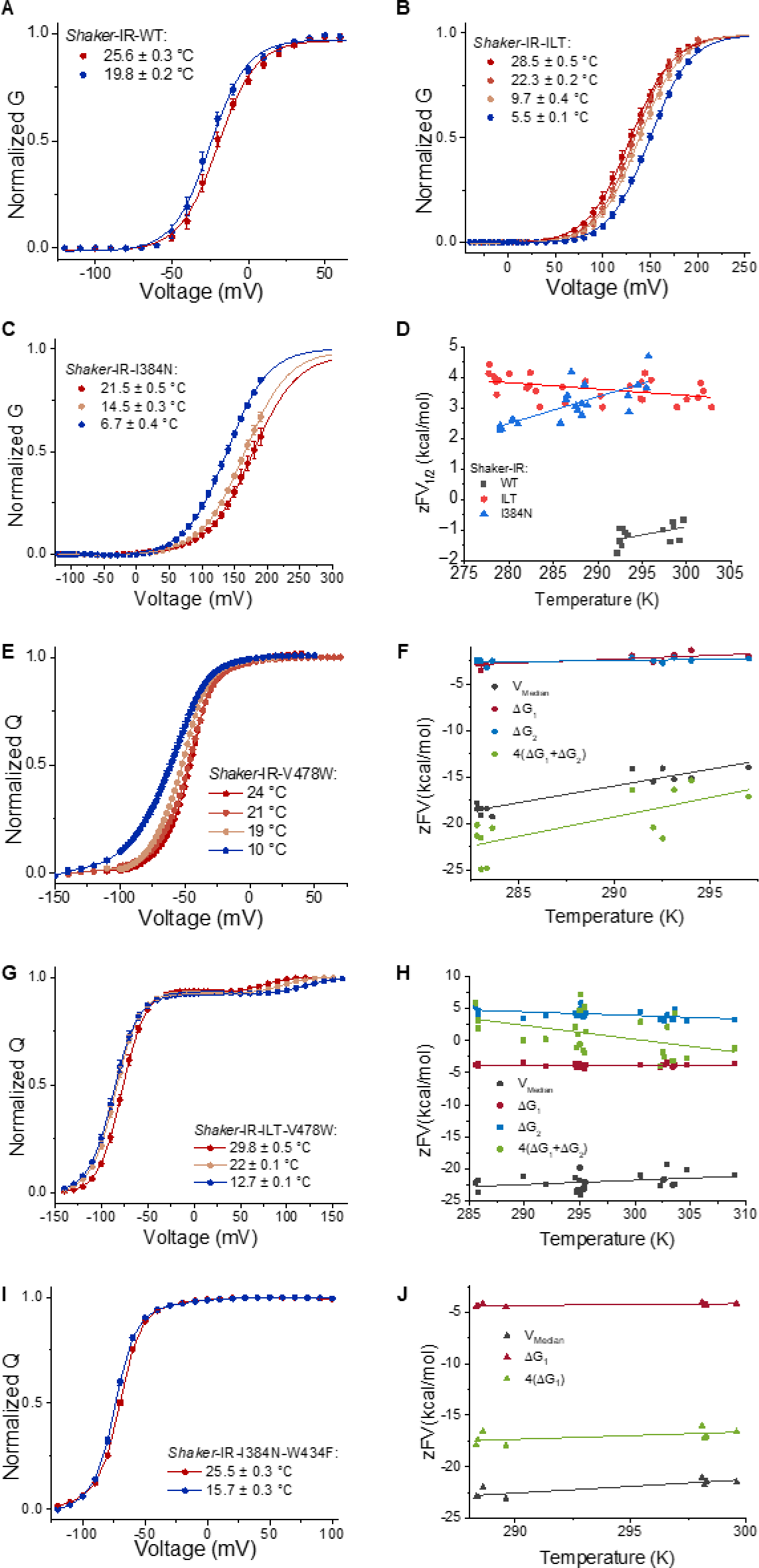
Thermodynamics of pore opening and VSD movement in WT, ILT, and I384N channels. (A-C) Normalized G-V plots for WT (A), ILT (B), and I384N (C) channels under various temperatures indicated by different colors. The continuous line corresponds to a two-state model fitting (parameters found in **Supplementary Table 1**). (D) Free energy vs temperature for the G-V relations. The WT (gray) and I384N (blue) constructs show a negative ΔS (-slope), while ILT (red) exhibits a positive ΔS. (E) Q-V relationship for the V478W channel (F) Free energy vs temperature plot for the V478W Q-V. (G) Q-V relationship for the ILT-V478W channel. (H) Free energy vs temperature plot for the ILT-V478W Q-V. (I) Q-V relationship for the W434F-I384N mutant. (J) Free energy vs temperature plot for the W434F-I384N mutant. The free energy vs temperature plot shows the energy for the first (ΔG_1_, red) and second (ΔG_2_, blue) component of the Q-V, the sum for the tetramer (4(ΔG_1_ + ΔG_2_), green) and using the V-median (gray). I384N only has one distinct component (ΔG_1_, red). The continuous lines in the Q-V curves are illustrative to help show the shifts.

The Q-V relationship for the WT (V478W) channel shows a displacement to the right with temperature **(Fig. 4E).** The difference between the Q-V curves for two temperatures is equivalent to the integral of the charge moved using the Tsteps (**Suppl. Fig. 5**). To analyze the thermodynamics of the WT Q-V curves, we used two complementary approaches: i) the Q-V was fitted using a three-state model where each transition has its own voltage dependence (Eq. 5); and ii) using the V-median, a model-independent procedure for calculating energies (25). We observe a good agreement in the WT between both approaches when we account for the contribution of all 4 voltage sensors **(Fig. 4F**, **Table 2)**. Unlike the V-median, the three-state model allows us to distinguish the contribution of each of the two main VSD movement transitions. With the fit to the three-state model, we find that the first component has the largest shift with temperature. Using the V-median fit, we found a shift of about 1.1 mV per °C that closely matches the G-V shift. This is expected due to the strict coupling between the VSD and PD in the WT channel, implying that both steps of gating charge movement contribute to the shift of the G-V in the WT channel.

**Table 2:** Thermodynamic parameters from Q-V curves.

|  | **WT** | **ILT** | **I384N** |
| --- | --- | --- | --- |
| **ΔH1 kcal/mol** | -26±6 | -5±2 | -10±3 |
| **ΔS1 cal/(mol\*K)** | -80±22 | -3±8 | -18±11 |
| **ΔV1/ΔT 1 (mV/°C)** | 2.3±0.3 | 0.9±0.1 | 0.31±0.02 |
| **ΔH2 kcal/mol** | -10±4 | 21±7 | - |
| **ΔS2 cal/(mol\*K)** | -24±15 | 58±22 | - |
| **ΔV2/ΔT (mV/°C)** | 0.5±0.1 | -2.6±0.2 | - |
| **ΔHT<sup>†</sup> kcal/mol** | -140±36 | 65±27 | -39±12 |
| **ΔST<sup>††</sup>cal/(mol\*K)** | -416±126 | 218±90 | -73±42 |
| **ΔHVmed kcal/mol** | -120±12 | -42±11 | -60±9 |
| **ΔSVmed cal/(mol\*K)** | -360±43 | -66±38 | -130±31 |
| **ΔVmed/ΔT (mV/°C)** | 1.1±0.1 | 0.46±0.06 | 0.43±0.03 |

<sup>†</sup> For WT & ILT 4(ΔH1+ ΔH2), for I384N 4(ΔH1)
<sup>††</sup>For WT & ILT 4(ΔS1+ ΔS2), for I384N 4(ΔS1)
Values shown as Mean±SE

In ILT case (ILT-V478W) the Q-V components are clearly distinct **(Fig 4G)**. We observe that the first component of the Q-V (80-90% of total charge) has a right shift of 0.9 mV/°C, while the second component (10-20% of total charge) has a left shift of -2.6 mV/°C. Quantifying the thermodynamics of ILT VSD movement shows a negative ΔS for the first transition, compared to a positive ΔS for the second transition (**Fig 4H**, **Table 2**). Using the V-median, we determine an overall shift of 0.46 m V/°C. This discrepancy can be explained by the fact that the V-median will be most affected by the first component of the Q-V that carries most of the charge. Thus, in this case, it provides less thermodynamic information of the energetics of the gating process. Our results for the WT and ILT indicate that the temperature dependence of the G-V arises from the temperature dependence of the VSD movement, and in the case of ILT it is dependent only on the second component. This is expected due to the separation of the Q-V components in ILT and the strict coupling between VSD activation and the opening of the channel, which has important implications for our understanding of channel function and temperature sensitivity.

A different picture emerges for I384N (W434F-I384N), where the Q-V shows only one component with a 0.31 mV/°C shift (**Fig 4I**, **Table 2**). The V-median, in this case, gives 0.43 mV/°C shift. The thermodynamic parameters for I384N Q-V show a reduced ΔS compared to the WT (**Fig 4J**, **Table 2**). The difference in the thermodynamic parameters between a two-state model fitting and the V-median procedure likely arises due to the underestimation of the energetics when fitting a multistep process with a simple two-step model (26). In this case, the V-median is expected to represent a better estimation of the thermodynamic parameters of the VSD movement. As indicated, the Q-V develops between -120 and -20 mV, voltages at which there is little or no opening of the channel (pore starts opening usually at -30mV for I384N), demonstrating that unlike in the WT channel and ILT, in this mutant, the shift of the G-V does not reflect the shift of the Q-V with temperature. Since this channel is not strictly VSD-PD coupled, this indicates that we are observing the temperature dependence of the pore itself.

## Discussion

Here, we used a newly developed technique to probe the temperature effects on the gating mechanisms of a Kv channel and two temperature-sensitive mutants. We have observed three major effects in the Shaker channel: i) the closure of the pore, which can be explained by the inward movement of the gating charges by a Tstep; ii) the direct movement of the gating charges in response to Tsteps; iii) mutations in the S4 and S4-S5 linker modulate the temperature dependence of the channel allowing us to determine the thermodynamics properties of specific transitions in the activation energy landscape. The effects of Tstep observed in the WT channel are in line with previous reports (5). The Tstep technique has the advantage over bath temperature changes in that it allows us to apply a fast temperature change at any given time during a voltage protocol, making it possible to reveal the characteristics of the energy landscape during channel gating. An example of this is the unprecedented observation of the movement of the gating charges by a temperature pulse, which led to the prediction of the temperature and time dependence of the Cole-Moore shift. This example demonstrates the potential of Tstep technique to dissect the energy landscape of ion channel gating mechanisms.

### Where does the temperature dependence of I384N and ILT come from?

We have shown that using Tstep, a temperature pulse closes the I384N channel, resembling the effect of the Tstep in TRPM8 observed previously (18). Therefore, the I384N channel mimics the response of a cold receptor. From previous kinetic analysis of the WT channel, we know that the last transition before channel opening has a shallow voltage dependence and large negative ΔH and ΔS components (-30 kcal/mol and -100 cal/K mol respectively) (6). This significant and negative enthalpic and entropic change would lead to a large rightward shift in the G-V curve with temperature, as observed in the I384N mutant. Since the I384N mutant effectively uncouples the movement of the VSD from the PD this mutation reveals the temperature dependence of the pore itself. Our data further shows that the pore opening is intrinsically affected by temperature. However, this is not observed in the WT channel. The WT channel does not change its G-V like I384N because of the strict coupling between VSD and PD. In this scenario, the individual energetic contribution of each step is combined to produce the temperature behavior of the G-V. Therefore, through detailed kinetics analysis of the WT channel or uncoupling the VSD-to-PD, we can observe this intrinsic property of the channel. In Shaker, the opening of the PD in one subunit is affected by the state of the PD gate in the adjacent subunit, giving rise to a highly cooperative opening of the PD. This highly cooperative opening requires activation of all the VSD thus the PD acts like a load on the VSD due to the strict coupling between VSD activation and PD opening. The uncoupling found in I384N suggests that the VSD in this mutant can move without the extra load of the PD, which would explain why we observe lower absolute values for ΔH and ΔS and why we only observe one component in the Q-V. Our mechanistic interpretation of the observed behavior of the I384N mutant is that the S4-S5 linker has a “loose” conformation, where movements of the VSD are not efficiently coupled to the S6 gate opening and temperature promotes the closure of the S6 gate.

Conversely, the ILT mutation had the opposite effect with temperature, increasing the amount of ionic current up to three-fold with a 5 °C Tstep, a response more in line with a heat receptor than a Kv channel. The ILT mutation introduces a voltage-dependent step that is also highly temperature-dependent and drives the concerted opening of the channel (16). It is not clear what step in the gating transition is the one affected by ILT, originally it was proposed to be the concerted opening of the pore, but later work using tetrameric concatemers of the channels has suggested that is the previous step, the last transition of the voltage sensor (15, 16). The ILT shares a similar effect as F290A mutation (27). Residue F290 determines the translocation of the last gating charge of the VSD necessary for the opening of the PD (28). The fact that mutations in F290 and ILT have additive effects in the G-V and last component of the Q-V support the notion that ILT also affects the translocation of the last charge, rather than the concerted opening step. By measuring the gating and ionic currents at high depolarized potentials, we observed a left shift in the G-V curve consistent with the shift in the second component of the Q-V. These shifts arise from the observed positive ΔS observed for the last transition in ILT (**Table 2**, **Eq. 6**). We believe this positive entropic change occurs at least in part due to conformational entropy of the VSD. We envision that in the intermediate state of the VSD in ILT (before the last gating charge moves), the S4 segment charges have less possible rotational conformation by the constraints imposed by the hydrophobic plug (29, 30). When the VSD moves from the intermediate to active conformation, the S4 charges are exposed to the solvent and thus have more conformational freedom, producing a net increase in the number of possible microstates, increasing entropy. This is not observed on WT because the last gating charge movement is lumped with the rest of the gating process, and we cannot distinguish the movement of the individual gating charges in that situation..Therefore in ILT the net change in entropy of the VSD would be the main driver of the Q-V and G-V shifts.

However, this hypothesis does not fully explain the observed increase in the ILT ionic current when the temperature increases. A clear example of this effect is when a Tstep is applied during a pulse to 180 mV, a voltage where the Q-V and G-V are saturated, but we still observed a three-fold increase in the current with a 5 °C step (**Fig. 1H**). Since ILT single-channel conductance is the same as the WT (16), it is unlikely that changes in the single-channel conductance can account for these effects. As mentioned above, it is possible that ILT also affects the last concerted step. At low temperatures, the S4-S5 linker in the ILT mutation also has a “loose” conformation where there is a fraction of the channels that do not open in response to the movement of the voltage sensor, and the S4-S5 linker is not able to propagate the electromechanical energy to the S6 gate. At high temperatures, the S4-S5 linker adopts a “tighter” conformation, propagating the motion of the VSD more efficiently to the S6 gate, opening a significant fraction of channels. This mechanism explains the increase in the Po and the increase in the rates of activation observed in the ILT mutant when Tstep is applied. The process of “loose” or “tight” coupling can be related to the interaction between the S4-S5 helix and the S6 C-terminus.

To illustrate how the proposed mechanism reproduce the observed effects of temperature, we built a kinetic model using the thermodynamic insight we have obtained (**Supplementary Figure 6 and Supplementary Table 2**). This model is based on Zagotta, Hoshi, Adrich model (14), where the voltage sensor moves in two steps (resting to intermediate and intermediate to active) and every voltage sensor must activate before the channel can open **(Figure 5A)**. Unlike the original model, we add an additional transition that represents the opening of the channel due to the opening of the bundle crossing gate (transition C15 to O – **Supplementary Figure 6A**). The transitions between these states are defined by the rates that describe the energy landscape of each mutant (**Supplementary Figure 6 B-D and Supplementary Table 2,3**). The voltage sensor transitions carry most of the charge movement, while the last opening transition has low voltage dependence. In the case of the WT, temperature affects mainly the resting to intermediate transition of the VSD by favoring the resting state and the last transition favoring the closed state consistent with our experimental results and previous determinations (6). The equilibrium of the last transition is displaced towards the closed states, but it does not produce a considerable change in the open probabilities by itself due to the strong coupling with the VSD. In response to a simulated Tstep this model produces an inward current due to the VSD movement that lead to a closing of the channel, at larger voltages where the Q-V saturates no such effect is observed: only an increase in current due to the increase in the single channel conductance is observed (**Figure 5B**). To reproduce the ILT phenotype, namely the split in the Q-V curves, we assume that the movement of most of the charge is carried in the resting to intermediate transition while a small fraction is accounted for in the intermediate to active transition (**Supplementary Table 2**). This accounts for the inferred effect of ILT in the translocation of the last gating charge. The ILT mutant exhibits a significantly larger difference in free energy in the intermediate to active transition with an enhanced temperature dependence compared to WT. Additionally, temperature favors the open state with temperature, reflecting the “tightening” of the coupling mechanism between the VSD and PD. In response to a Tstep this model produces a biphasic effect on the gating currents and opening of the channel (**Figure 5C**). On the other hand, in I384N only one transition of the VSD was observed so we included most of the ΔH and ΔS in the first transition of the VSD. In this case, the opening of the channel exhibits a shallow voltage dependence, and the ionic current movement is displaced compared to the VSD movement. The large negative ΔS would produce a large rightward shift in the voltage dependence with temperature, promoting closure. In response to a Tstep this model produces an inward current due to the VSD movement like the WT and a closing of the channel at all voltages tested (up to +150 mV) (**Figure 5D**). Our model can provide a visual and thermodynamic representation which aids in understanding the experimental observations and proposed gating mechanisms for the WT and mutant channels (**Figure 1, 2, 3 and 4)**. These models reproduce the observed properties of the Q-V, G-V and temperature displacement of the gating currents for the WT, ILT and I384N (**Figure 5E-G**). Overall, this modeling effort further demonstrates that changes in the energetics of VSD-PD coupling can turn a prototypical Kv channel into a temperature receptor.

**Figure 5:**
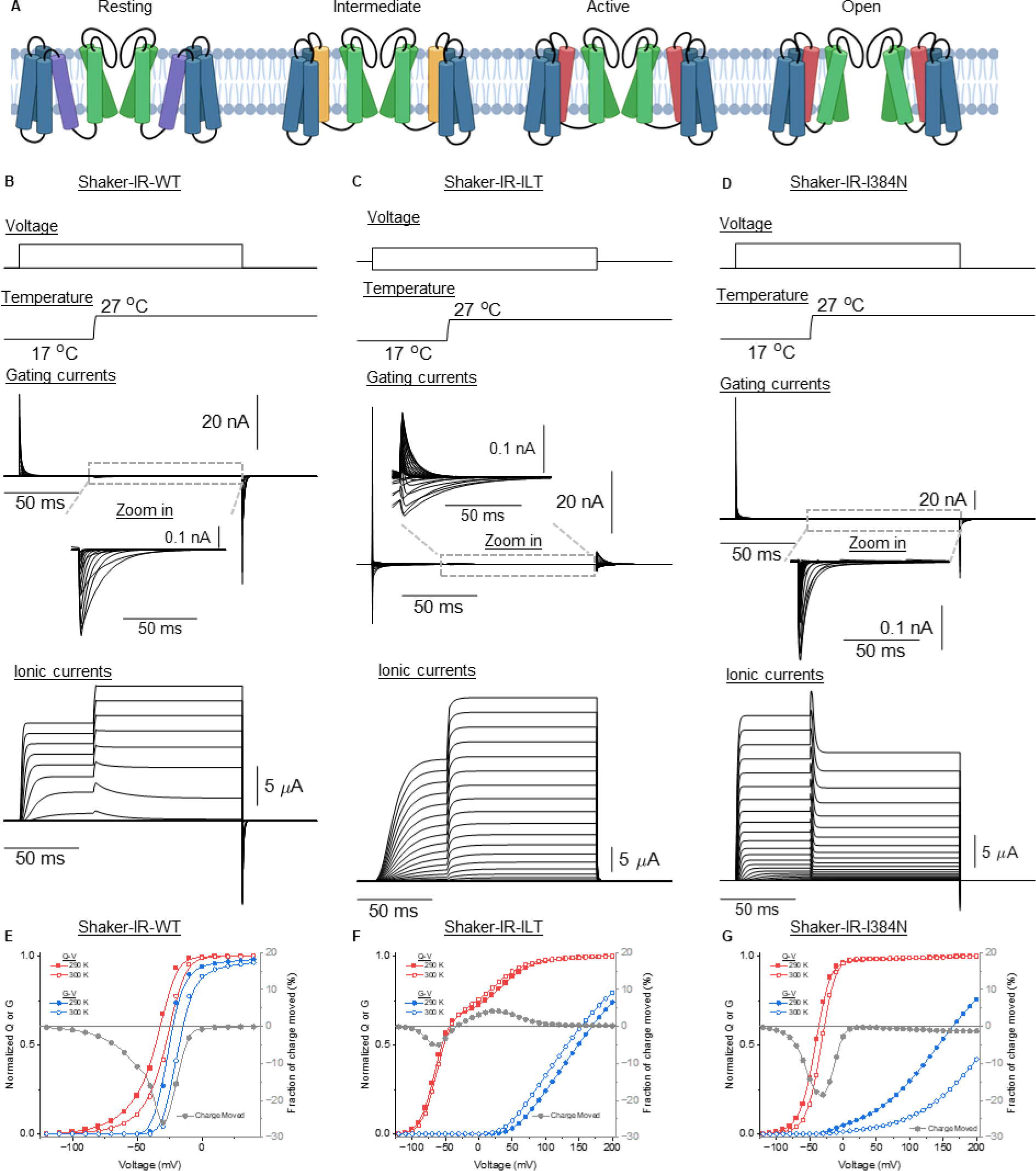
Simulating the effects of temperature steps on the gating transitions of *Shaker* WT, I384N and ILT. (**A**) Schematic representation of different conformational states of the channel, from left to right: Resting (VSD down, pore closed), Intermediate (VSD intermediate, pore closed), Active (VSD up, pore closed), and Open (VSD up and pore open). The color of the transmembrane helix denotes the movement of the S4 segment in the resting (purple), intermediate (yellow) and active state (red). (**B**), (**C**), and (**D**) are the simulations of the ionic and the gating currents when Tstep is applied at the middle of a voltage protocol. The model used and equations are shown in **Supplementary figure 6**. The parameters used for simulation are shown in the **Supplementary Table 2 & 3**. **(E-G)** Simulated Q-V (red), G-V (blue) and the fraction of the charge moved (gray) for the WT(**E**), ILT (**F**) and I384N (**G**) mutants at 290 (filled symbols) and 300K (empty symbols). The voltage protocols used are the same as presented in Figure 1.

Our mechanistic interpretations suggest that the S4-S5 linker can adopt either a “loose” or “tight” conformation, depending on the mutations and temperature conditions. In the “loose” conformation, the movement of the VSD is necessary but not sufficient to efficiently propagate the electromechanical energy to the S6 gate, leading to a reduced open probability. Conversely, in the “tight” conformation, the S4-S5 linker more effectively couples the VSD motion to the S6 gate, opening the PD. This functional switching is related to the coupling mechanism and can be tuned by interactions between the intracellular part of the S4, S4-S5 linker and the C-terminus of the S6 segment. Our results show that temperature dependence can arise by changes in the coupling between VSD and PD.

The Shaker K channel has a strict coupling of the voltage sensor with the pore opening. This contrasts with thermo-TRP channels, where the coupling of the voltage and temperature sensors and the pore is allosteric (31). The present study has given possible molecular mechanisms for transforming the channel into cold or heat receptors by introducing mutations that can change the magnitude of the VSD-pore coupling at different positions along the Shaker channel activation energy landscape. However, they should not be interpreted as equivalent to the mechanisms of the thermo-TRP channels. Instead, the present study shows how a strictly coupled channel can be converted from heat into a cold receptor. Still, most importantly, it has allowed a dissection of the energy landscape of the voltage sensor movement and its coupling to the opening of the pore.

## Materials and methods

### Channels expression in *Xenopus* oocytes

*Xenopus laevis* ovaries were purchased from Xenopus 1 (Dexter, Michigan). The follicular membrane was digested by 2 mg/ml collagenase supplemented with bovine serum albumin (BSA) 1mg/ml. Oocytes were kept at 12 or 18 °C in SOS solution containing (in mM) 96 NaCl, 2 KCl, 1 MgCl_2_, 1.8 CaCl_2_, 10 HEPES, pH 7.4 (NaOH) supplemented with gentamicin (50 mg/ml). After 6 - 24 hr of harvesting, they were injected with 5-50 ng of cRNA diluted in 50 nl of RNAse-free water and incubated for 1-4 days before recording. We used clones from Shaker zH4 K_+_ channel with removed N-type inactivation (IR, Δ6-46) in the pBSTA vector (32). Mutations were performed using Quick-change site-directed mutagenesis and cRNA was transcribed from linearized cDNA, using mmessage mmachine T7 Transcription kit from Invitrogen (Waltham, Massachusetts). cDNAs were sequenced to attest to the correct sequence.

### Electrophysiology

Ionic and gating currents were recorded from oocytes using the cut-open voltage-clamp method (33). Voltage-sensing pipettes were pulled using a horizontal puller (P-87 Model, Sutter Instruments, Novato, CA), and the resistance ranged between 0.2-0.5 MΩ. Data were filtered online at 20–50 kHz using a built-in low-pass four-pole Bessel filter in the voltage clamp amplifier (CA-1B, Dagan Corporation, Minneapolis, MN, USA) sampled at 1 MHz, digitized at 16-bits and digitally filtered at Nyquist frequency (USB-1604; Measurement Computing, Norton, MA). The voltage command and the current elicited were filtered using the same frequency. An in-house software was used to acquire (GPatch64MC) and analyze (Analysis) the data. The chamber temperature was measured by a thermocouple and controlled through a negative feedback loop using a Peltier cooler. Transient capacitive currents were subtracted from the recorded currents by a dedicated circuit. We used an offline linear subtraction procedure to remove the optocapacitive and leak effects of Tsteps using voltages at which there is no ionic current or gating charge movement (18). For ionic current measurements, the external solution was composed of (mM): KOH 12, CaOH_2_ 2, HEPES 10, EDTA 0.1, N-methyl-D-glucamine (NMDG) 108 and the internal solution of (mM): KOH 120, EGTA 2, HEPES 10. For gating current measurements, the external solution was composed of (mM): NMDG 120, CaOH_2_ 2, HEPES 10 EDTA 0.1 and the internal solution of: NMDG 120, EGTA 2, HEPES 10. In all cases, all the solutions were adjusted to pH 7.4 with methanosulfonic acid. For I384N-W434F mutant oocytes were incubated for 30-60 min in an internal solution prior to the experiment to dialyze the internal potassium. All chemicals used were purchased from Sigma-Aldrich (St. Louis, MO).

### Tstep set-up

A 3.5 W 447nm diode laser (Osram PLPT9 450D_E A01) was placed on top of the recording chamber and aligned with the oocyte dome. The beam was collimated using an aspheric lens (A230TM-A,Thorlabs) followed by a pair of microlens arrays (MLA300-14AR-M, Thorlabs Inc. Newton, New Jersey) to ensure homogenization of the beam (34), and focused into the oocyte dome using another aspheric lens (ACL2520U-A,Thorlabs Inc. Newton, New Jersey). An in-house current modulated power supply was used to achieve rapid turn-on of the laser. The laser was triggered by an arbitrary wave generator (4075B, B&K Precision corp., Yorba Linda, CA).

### Capacitance-based temperature measurement (CTM)

We measured the temperature due to Tstep as reported previously (18). Briefly, we applied a sinusoidal voltage wave during the Tstep. We calculated the complex voltage and current signal after subtraction of the optocapacitive current using the Hilbert transform. We calculate the impedance by dividing the complex voltage and current. The impedance of the system is given by:

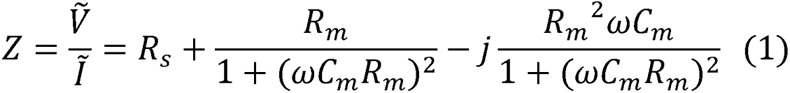

where Rs, Cm, Rm, ω and j are the series resistance, the membrane capacitance, the membrane resistance, the angular frequency of sinusoidal wave and the imaginary unit, respectively. When (ω*C_m_R_m_*)^2^>>>1 we get

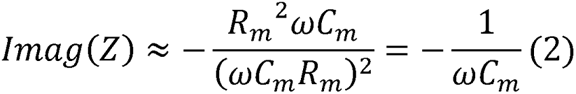

Thus, we can obtain the membrane capacitance time course using the time course of the imaginary part of the impedance.

We used the average of 100 traces and filtered the results using a 10KHz *offline* Bessel-like filter to obtain the capacitance time course. A linear fitting that converts capacitance to temperature was used to get the temperature (18). To ensure that the assumption of Eq 2 is valid, we worked at voltages in which there is no conductance for the conductive channels or no gating movement for the non-conductive channels.

### Data analysis

The temperature-dependent (Q_10_) factor for the ionic currents was obtained by:

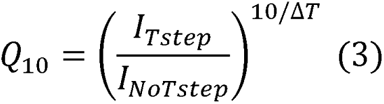

where I_Tstep_, I_NoTstep_, and ΔT correspond to the current at the end of the Tstep, the current right before the Tstep is applied and the temperature difference of the Tstep, respectively.

The G-V curves were measured from the tail currents after a voltage protocol and fitted using a two-state model given by equation:

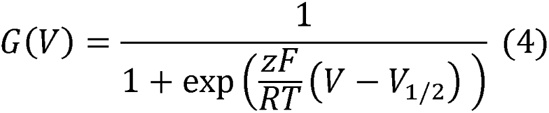

Where z is the apparent charge expressed in units of elementary charge (*e*_0_), V is the voltage and *V*_½_ is the voltage of half maximal conductance. R, T and F have their usual meanings.

For the analysis of the Q-V curves we used three different approaches:

1-A two-state model fitting equivalent to the one in Equation 4 was used to fit the W434F_I384N mutant and the individual components of the ILT Q-V.

2-A three-state model fitting is given by the following equation (35):

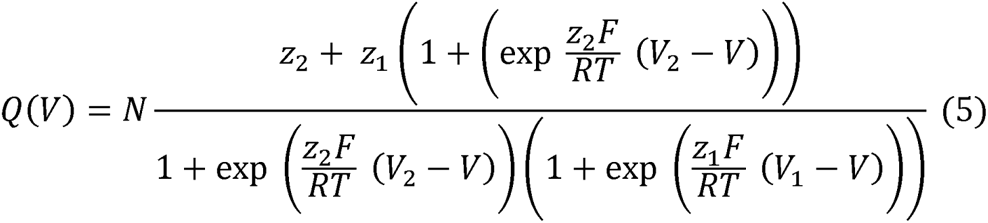

where N, z_1_, z_2_, V_1_ and V_2_ are the number of channels, the charges associated and equilibrium voltages for the first and second transition respectively. For the fitting at different temperatures is assumed that N, z_1_ and z_2_ remain constant.

3-Using the V-median of the Q-V (25). For the V-median the that number of charges per channel (z) was considered the same for all channels: 13.6 elementary charges (12, 36, 37)

To obtain the ΔS and ΔH of the G-V and Q-V curves we used the following linear relationship (38):

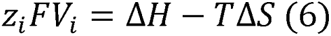

where the z_i_ and V_i_ represent the relative charges and voltages obtained from the fitting of the G-V and Q-V curves.

## Author contributions

Performed and analyzed experiments: CB, BP

Interpreted results: CB, BP, RL, FB

Conceptualization: CB, BP, RL, FB

Writing: CB, BP, RL, FB

Supervision: RL, FB

## Funding

National Institutes of Health Award R01GM030376 (FB, RL)

Fondo Nacional de Desarrollo Cientıfico y Tecnologico (FONDECYT). Regular Grant Number 1230267 ((RL)

PEW Latin American Fellow 2019 (BP)

## Acknowledgments

CB and BP agree that both authors should be the first author and the order of appearance was determined by coin toss.

## Code availability

Code for CTM analysis is freely available at github via: https://github.com/PintoBI/Tjumps under MIT license.

