## supplementary info for "Turning a Kv channel into hot and cold receptor by perturbing its electromechanical coupling"

**Correspondence**

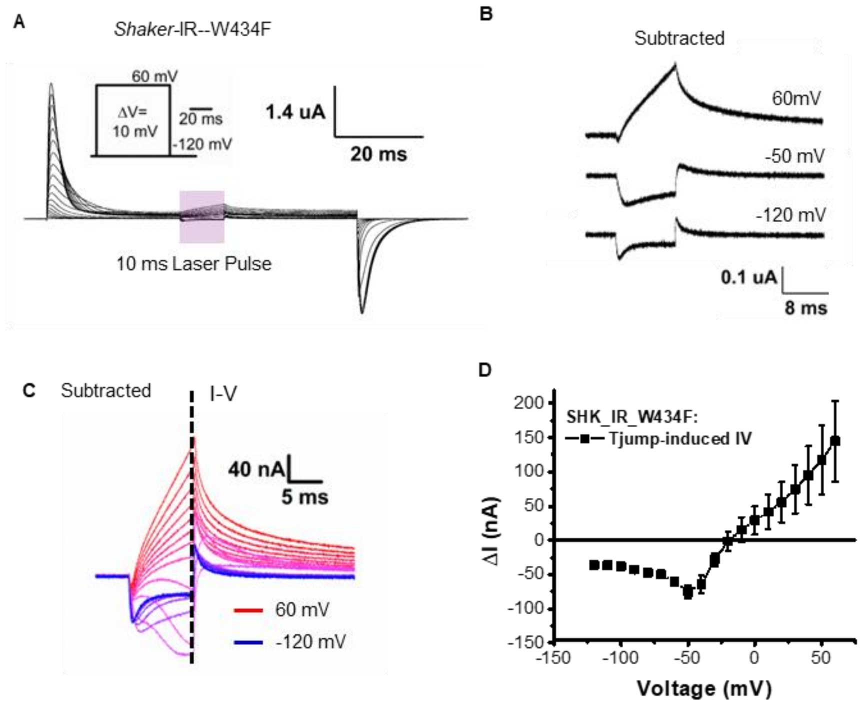

**Supplementary Figure 1: Effects of temperature into the W434F mutant. (A)** Gating current measurements during a voltage and laser step protocol for W434F. A temperature change was produced by a continuous laser pulse applied in the middle of a voltage step, after gating currents had subsided. **(B)** Detail into representative current traces following the subtraction of the linear optocapacitive component showing the appearance of ionic currents at positive voltages. **(C)** Current traces after the subtraction of the linear optocapacitive component the line indicates the isochronal used to measure the current to voltage (I-V) relationship. **(D)** I-V curve after subtraction showing a positive current due to the appearance of ionic conduction.

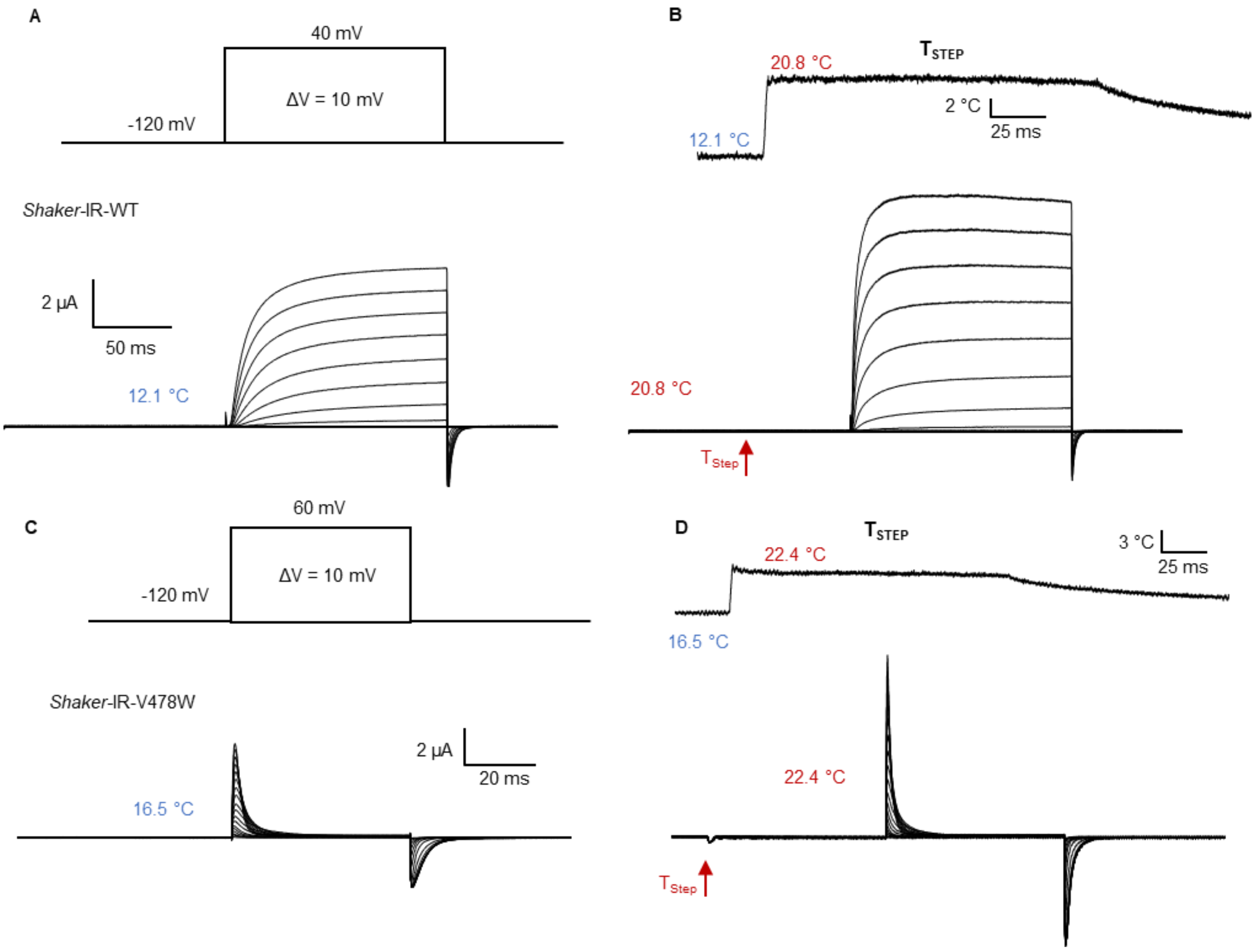

**Supplementary Figure 2: Representative traces for G-V and Q-V measurements in WT channel. (A,B)** Ionic currents in response to a voltage pulse protocol for Shaker-IR in the absence **(A)** and presence **(B)** of a Tstep before the voltage step. **(C,D)** Gating currents in response to a voltage pulse protocol for Shaker-IR-V478W in the absence **(A)** and presence **(B)** of a Tstep before the voltage step. The Voltage step protocol and Tstep profile is shown over the recordings. Bath temperature shown in blue, Tstep temperature shown in red, arrows show timing of Tstep application.

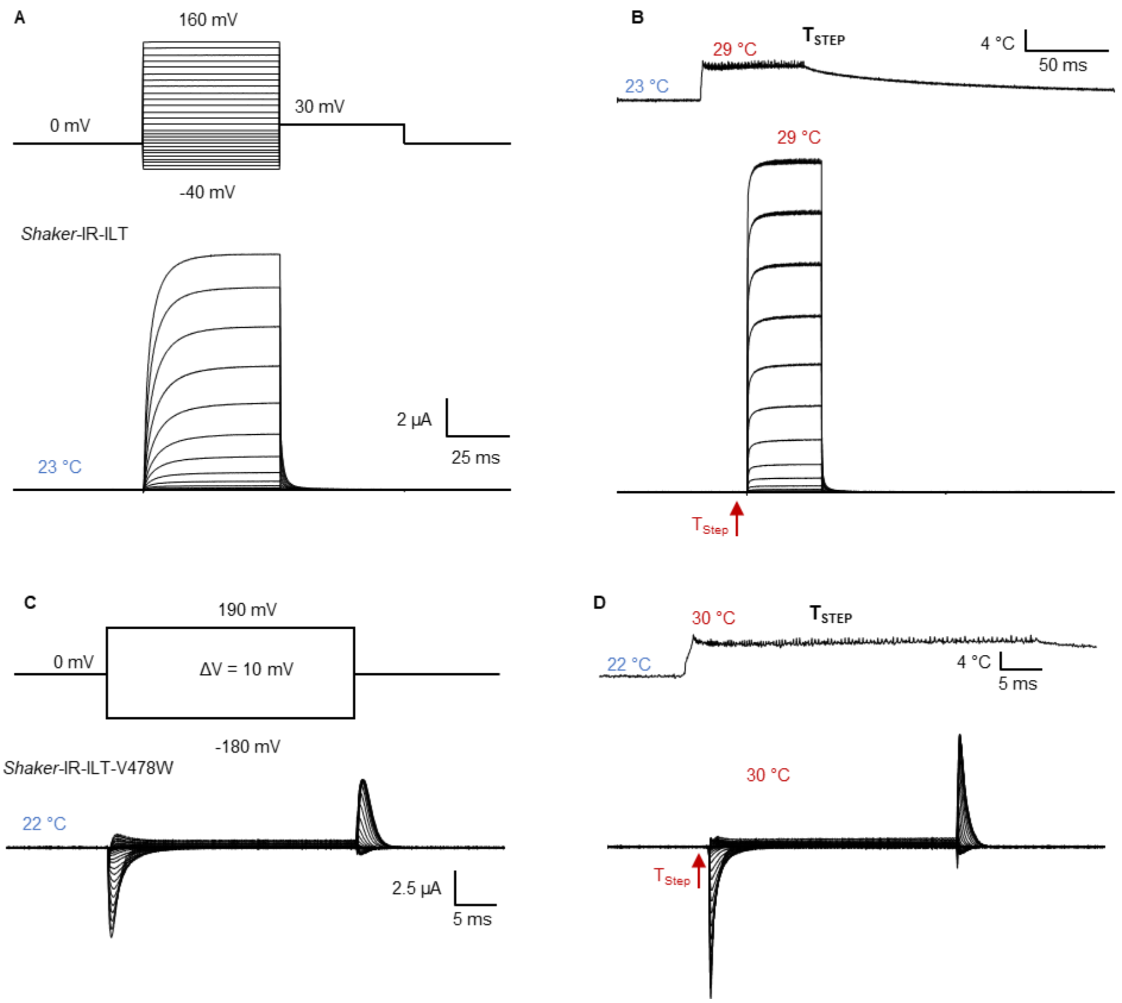

**Supplementary Figure 3: Representative traces for G-V and Q-V measurements in ILT mutant. (A,B)** Ionic currents in response to a voltage pulse protocol for Shaker-IR-ILT in the absence **(A)** and presence **(B)** of a Tstep before the voltage step. **(C,D)** Gating currents in response to a voltage pulse protocol for Shaker-IR-ILT-V478W in the absence **(A)** and presence **(B)** of a Tstep before the voltage step. The Voltage step protocol and Tstep profile is shown over the recordings. Bath temperature shown in blue, Tstep temperature shown in red, arrows show timing of Tstep application.

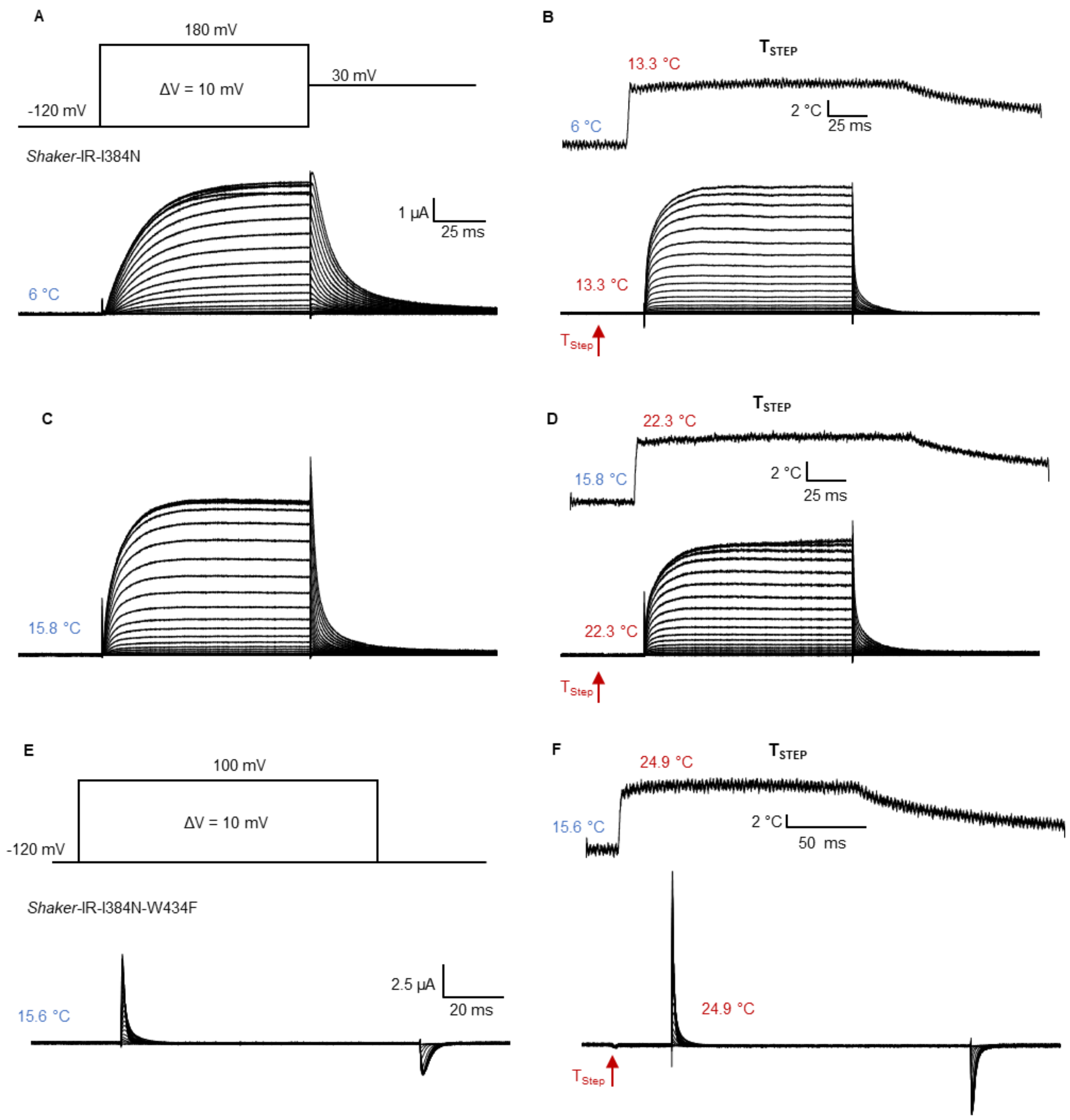

**Supplementary Figure 4: Representative traces for G-V and Q-V measurements in I384N mutant. (A,B)** Ionic currents in response to a voltage pulse protocol for Shaker-IR-I384N in the absence **(A)** and presence **(B)** of a Tstep before the voltage step. **(C,D)** Gating currents in response to a voltage pulse protocol for Shaker-IR-I384N-W434F in the absence **(A)** and presence **(B)** of a Tstep before the voltage step. The Voltage step protocol and Tstep profile is shown over the recordings. Bath temperature shown in blue, Tstep temperature shown in red, arrows show timing of Tstep application.

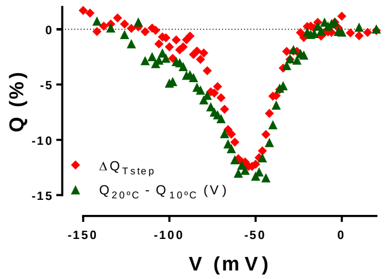

**Supplementary Figure 5: Charge moved by temperature vs voltage.** Fraction of the total gating charge moved at different voltages for a 10º C Tstep from a bath temperature at 10º C. The charge was calculated by integration of the Tstep induced current (red) and by subtraction of the Q-V curves at different temperatures (green).

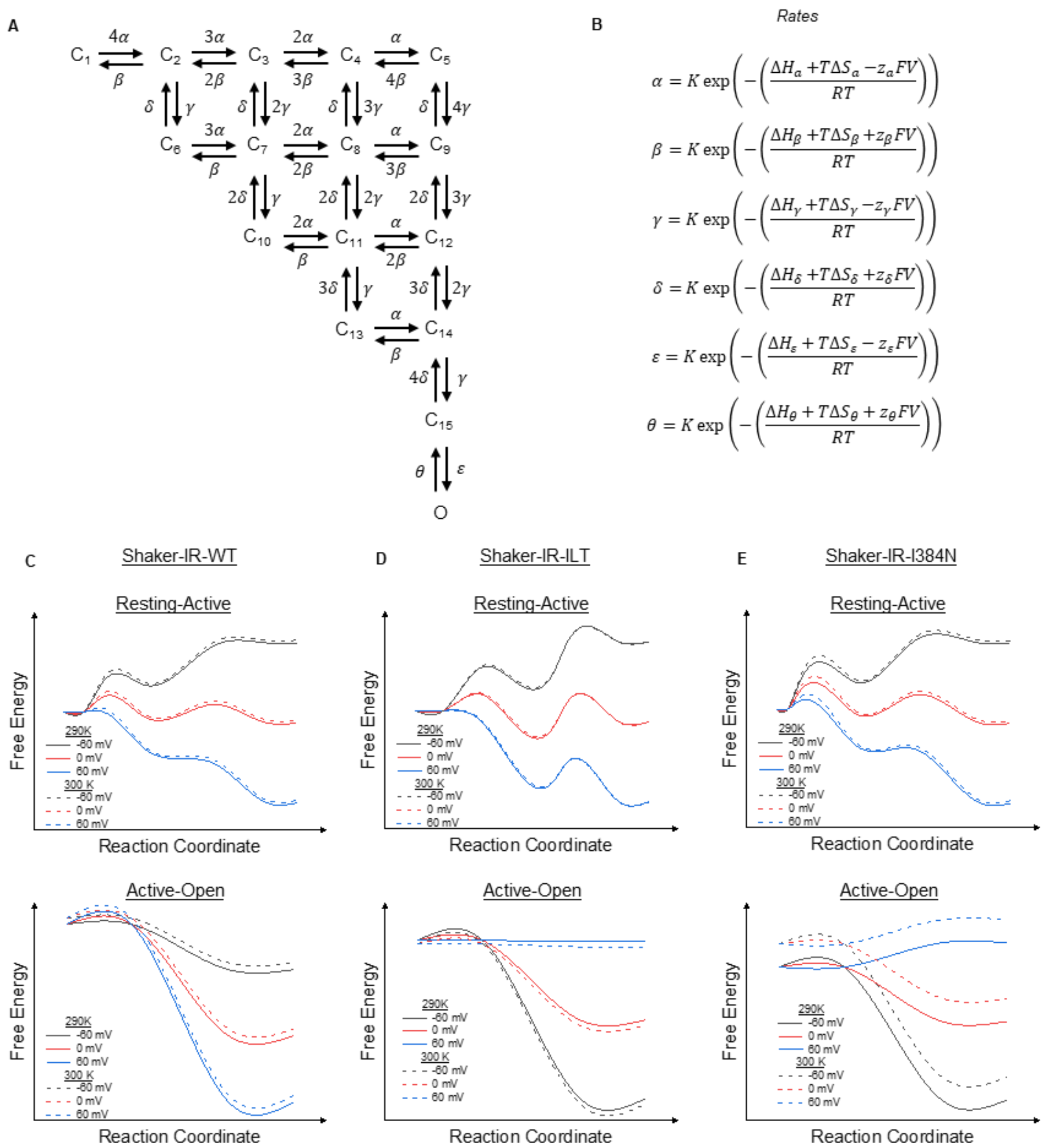

**Supplementary Figure 6: Kinetic model for Shaker-IR channels used for simulating ionic and gating currents. (A)** Kinetic model based on Zagotta, Hoshi and Aldrich model. The model consists of 16 states, where 15 are closed states and one open state. The pore only opens when all the voltage sensors activate. The transition from state 15 to O represents the open of the bundle crossing in the pore. **(B)** Rates constant used for the model in A. The values used for the enthalpic, entropic components and the charge associate with each transition is shown in the supplementary table 2. K is a constant related to the vibration frequency. This factor is linearly dependent on temperature and has the same value for all the transitions (500/ms at 290K). **(C-E)** Energetic representation of the transitions between Resting to Active (upper) and Active to Open (lower) state for WT **(C)**, ILT **(D)** and I384N **(E)** at -60 (gray), 0 (red) and 60 (blue) mV for temperature 290 (solid lines) and 300K (dashed lines). The curves where calculated using the parameters in **Supplementary table 2, 3**.

**Supplementary Table 1: Fitting parameters for figure 4A-C**

| | z | V1/2 |
|---|---|---|
| **WT** | | |
| **25.6ºC** | 2.4±0.1 | -18.7±0.7 |
| **19.8ºC** | 2.6±0.1 | -23.3±0.7 |
| **ILT** | | |
| **28.5ºC** | 1.41±0.02 | 130±1 |
| **22.3ºC** | 1.57±0.02 | 134±2 |
| **9.7ºC** | 1.57±0.02 | 138±2 |
| **5.5ºC** | 1.62±0.02 | 150±2 |
| **I384N** | | |
| **21.5ºC** | 0.95±0.01 | 179.0±0.4 |
| **14.5ºC** | 1.01±0.01 | 166.0±0.3 |
| **6.7ºC** | 1.108±0.008 | 138.60±0.2 |

Values shown as Mean±SE

**Supplementary Table 2: Steady state parameters used for simulating WT, ILT and I384N channels.**

| **Resting to intermediate** | **WT** | **ILT** | **I384N** |
|---|---|---|---|
| **ΔH kcal/mol** | -18.8 | -9.5 | -18.8 |
| **ΔS cal/(mol*K)** | -60 | -20 | -60 |
| **z** | 1.4 | 2.4 | 1.4 |
| **Intermediate to active** | | | |
| **ΔH kcal/mol** | -3.8 | 9.8 | -4.5 |
| **ΔS cal/(mol*K)** | -8 | 31 | -8 |
| **z** | 2.1 | 1.1 | 2.1 |
| **Active to open** | | | |
| **ΔH kcal/mol** | -10.5 | 7.5 | -23 |
| **ΔS cal/(mol*K)** | -30 | 20 | -85 |
| **z** | 0.5 | 0.5 | 0.5 |

**Supplementary Table 3: Parameters used for simulating WT, ILT and I384N channels.**

| WT | | | |
|---|---|---|---|
| **Transition** | **ΔH (kcal/mol)** | **ΔS (cal/mol)** | **z (e)** |
| α | -22.8 | -92 | 0.7 |
| β | -4 | -32 | 0.7 |
| γ | -5.8 | -31 | 1.2 |
| δ | -2 | -23 | 0.9 |
| ε | -4.75 | -27 | 0 |
| θ | 5.75 | 3 | 0.5 |

| | | | |
|---|---|---|---|
| ILT | | | |
| **Transition** | **ΔH (kcal/mol)** | **ΔS (cal/mol)** | **z (e)** |
| α | -6 | -35 | 1.1 |
| β | 3.5 | -15 | 1.3 |
| γ | 11 | 19 | 0.2 |
| δ | 1.2 | -12 | 0.9 |
| ε | 8.4 | 13 | 0 |
| θ | 0.9 | -7 | 0.5 |
| I384N | | | |
| **Transition** | **ΔH (kcal/mol)** | **ΔS (cal/mol)** | **z (e)** |
| α | -21.8 | -90.2 | 0.4 |
| β | -3 | -30.2 | 1 |
| γ | -1.5 | -22.7 | 1 |
| δ | 3 | -14.7 | 1.1 |
| ε | -21 | -85 | 0 |
| θ | 2 | 0 | 0.5 |
